# Exploring *Porphyromonas gingivalis* Pathogenesis and Antibody Pool Effect Using a Novel Mouse Reproductive Model

**DOI:** 10.1101/2024.06.28.601233

**Authors:** Xiaoju Shi, Maria Xu

**Affiliations:** Department of Research and Development, Yangtze Green Biotechnology Co., Ltd. CN; Manchester Centre for Genomic Medicine, UK

**Keywords:** *P. gingivalis*, pathogenesis, outer membrane proteins, antibody pool effect, mouse reproductive model

## Abstract

This study introduces a novel animal model consisting of two phases. In phase I, Balb/c mice received three intraperitoneal injections of antibodies and two subcutaneous challenges with *Porphyromonas gingivalis*. The mice were divided into seven groups: G1, G2, and G3 received a single monoclonal antibody targeting the major outer membrane protein RagB of *P. gingivalis* and infection; G4 was infected without antibody; G5 and G6 received a mixture of monoclonal antibodies and infection; and G7 served as the healthy control. In phase II, one male was paired with one female from the same group and one healthy female. Fertility was assessed using maximum body weight growth rate (MWGR) and alive pup rate (APR).

Results indicated that the APR of neonatal mice in G5+G6 significantly lower than G1-G3 and G7 (P < 0.05). Similarly, the MWGR of G5+G6 significantly lower than G7 (P < 0.05). Notably, the APR and MWGR in G4, which received no antibody, were lower than in G7 but higher than in G5-G6. These findings suggest that a mixture of antibodies coexisting with *P. gingivalis* infection leads to more detrimental reproductive damage than infection alone. The so-called Antibody Pool Effect plays a role in the pathogenesis of *P. gingivalis*. Understanding the mechanisms of this immunopathological damage may provide insights into the correlation between *P. gingivalis* infection and systemic diseases.

## INTRODUCTION

*P. gingivalis* has been recognised as the major causative agent of periodontal diseases since the early 1980s (1-3). Periodontal disease progresses through stages of acute infection, remission, recurrence, and eventually chronic persistent inflammation. Both clinical and animal studies have shown that *P. gingivalis* infection can damage local tissues.

Clinical treatment of periodontal disease typically involves broad-spectrum antibiotics, physical or mechanical removal of local lesions, and surgical interventions when necessary. However, *P. gingivalis* infections are often not completely eradicated. Research has focused on the prevention, treatment, and control of refractory *P. gingivalis* infections, including the development of biological products like vaccines and antibodies (4-11). To date, no curative products are available on the market. Bacterial genetic polymorphisms are an important factor in the development of immune protection products (7, 9, 11-13). Furthermore, due to the unclear pathogenic mechanisms, there are limited standards by which to measure the effectiveness of a treatment, and the endpoint evaluation of new drugs.

Conventional *P. gingivalis* infection animal models mainly focus on soft tissue destruction, alveolar bone resorption, and serum inflammatory index detection (4, 7-9, 11, 14-16). In these models, local tissue damage is observed in almost all animals, depending on the infection dose. Generally, the higher the bacterial concentration, the greater the local tissue damage. In the soft tissue destruction model, infected animals typically heal spontaneously, though some abscesses may reappear after initial wound healing (9). Mortality rates are usually low when animals are infected with *P. gingivalis* alone.

*P. gingivalis* infection is not only responsible for oral periodontal diseases but is also linked to various chronic systemic conditions such as cardiovascular disease (17-19), Alzheimer’s disease (20-23), diabetes, renal disease, gastrointestinal and respiratory system tumours (24-30), and adverse pregnancy outcomes (31-37), including low birth weight and premature births. The connection between local infection and systemic disease has previously been unclear, and experimental results sometimes conflict with proposed hypotheses.

Inspired by clinical findings, we established a mouse reproductive model to investigate the impact of *P. gingivalis* infection on reproduction. In our first experiment, we designed three offspring groups: (T1) both male and female mice received a combination of rabbit polyclonal antibodies and *P. gingivalis* infection; (T2) healthy male and female mice received a combination of antibodies and *P. gingivalis* infection; (T3) male mice received antibodies and *P. gingivalis* infection and were paired with healthy females.

Our results showed that in T1, where both parents were injected with antibodies followed by *P. gingivalis* infection, there was a low rate of live births and a high rate of stillbirths (15 live and 6 stillborn pups from 12 females in total). In T2, 12 females produced 51 live pups and 2 stillbirths. In T3, 12 females produced 22 live pups and 5 stillbirths (11).

This was our first experiment involving mouse breeding with *P. gingivalis* infection, and we recognised the need for improved experience and preparation. The data indicated that the infection status of males, with coexisting antibodies and *P. gingivalis* infection, had a greater impact on live birth rates than that of females.

Most scientific studies on pregnancy abnormalities associated with periodontal disease or *P. gingivalis* infection have focused on the maternal reproductive system and foetal characteristics, including genetics, endocrine characteristics, and tissue and organ infection (32-38). Our results suggest that paternal infectious status plays a crucial role in pregnancy abnormalities. We conducted further experiments to investigate the effects of single antibodies and antibody mixtures on reproduction. The aggregate data support our hypothesis that immune complexes of antigens and polyclonal antibodies have the potential to cause adverse pregnancy outcomes.

## MATERIALS AND METHODS

***P. gingivalis* CULTURE**. *P. gingivalis*, consisting of various subtypes, was grown on Fastidious Anaerobic Agar (FAA) plates containing 5% defibrinated horse blood. The plates were placed in an anaerobic incubator bag with an anaerobic gas-generating reagent (Anaero, Japan) at 37°C for at least 48 hours. The bacterial colonies on the plate surface were transferred to pre-warmed Brain Heart Infusion (BHI) medium containing 5 µg/ml Hemin and 10 µg/ml Vitamin K3. The cultures were grown for 48 hours under anaerobic conditions, then diluted 1:10 in fresh BHI culture medium and incubated for 16-18 hours until the OD540nm value reached 1-1.2. The culture was centrifuged and washed twice in PBS (pH 7.4) to prepare a bacterial suspension at an appropriate concentration (2-8 × 10^10^ CFU/ml) in PBS (pH 7.4).

### ANTIBODY RESOURCES

#### Polyclonal Antibodies

Mouse and rabbit antibody sera were prepared by immunising animals with antigenic protein three or four times, based on their antibody response. The titre of specific antibodies in sera was determined by indirect ELISA. Once the antibody level reached a set standard, whole blood was collected from the animal, and the serum was obtained after incubation and centrifugation, then frozen for storage.

#### Monoclonal Antibodies

The monoclonal antibodies against *P. gingivalis* outer membrane proteins were prepared by a third-party Contract of Research Organisation (CRO, HuaBio, CN). The general process of obtaining a monoclonal antibody involved immunising animals with antigenic proteins five times, with each immunisation interval being 1-2 weeks. Murine splenocytes were fused with well-grown SP2/0 myeloma cells and grown for seven days after fusion. Positive clones were screened with immunisation antigens and further subcloned three times to ensure stable antibody production. The amplified hybridoma seed cell lines were stored in liquid nitrogen.

### ANIMAL MODELS

#### Animal Selection

The experimental design used 56 male and 112 female Balb/c mice, purchased from Weitonglihua Technology Co., Ltd. (Beijing, CN). The mice were 9 to 10 weeks old and weighed 18 to 20 g.

#### Animal Feeding and Management

Experimental animals were housed in SPF-grade barrier facilities at the animal centre of Hongren Biopharm (Wuhan, CN). Before the start of the experiment, the animals underwent an isolation and adaptation period of 3-7 days. Environmental parameters were controlled, including temperature (20-26°C), humidity (40-70%), ventilation (10-20 times/hour), and a 12-hour light/dark cycle (lights from 7 a.m. to 7 p.m.; dark from 7 p.m. to 7 a.m.). The animals were continuously supplied with cobalt-60 radiation-sterilised forage and water.

#### Animal Ethics

This experiment was reviewed and approved by the Institutional Animal Care and Use Committee of Hongren Biopharmaceutical Co. Ltd (No. IACUC202300053, CN). The study complies with the relevant laws and regulations on the use and management of experimental animals. This study complies with the relevant laws and regulations on the use and management of experimental animals. The entire experimental process was completed in a single-blind manner at the Hongren Animal Centre Research Institute.

#### Grouping

56 male mice and 56 female mice were equally divided into 7 groups, with 8 mice in each group. Another 56 healthy control female mice were included in the study.

#### Medicine Administration

On the 2nd, 5th, and 21st days after grouping, animals in groups G1 to G6 were injected intraperitoneally with antibodies according to the schedule in Table 1.

**TABLE 1.** Animal experiment design and operation arrangement.

| Groups | Monoclonal antibody injection on Day 2, Day 5 and Day 21 | *P. gingivalis* infection on Day 6 and Day 22 | Male | Female |
| --- | --- | --- | --- | --- |
| G1 | 1D2-2-1-3 | yes | 8 | 8 |
| G2 | 1B11-4-4 | yes | 8 | 8 |
| G3 | 1C3-7-8 | yes | 8 | 8 |
| G4 | N/A | yes | 8 | 8 |
| G5 | 1B11-4-4+1C3-7-8 | yes | 8 | 8 |
| G6 | 1D2-2-1-3+1B11-4-4 | yes | 8 | 8 |
| G7 | N/A | N/A | 8 | 64 |

#### Murine Soft Tissue Destruction Model

A bacterial suspension was prepared at a concentration of 3×10^10^ CFU/ml. On the 6th and 22nd days after grouping, animals in groups G1 to G6 were challenged with *P. gingivalis* (strain 01-033, Bristol, *P. gingivalis* RagB-3 subtype) subcutaneously in the middle of the scapula on the back of the mice, 200 µl/mouse. From the beginning of the challenge to the peak of damage, the development of any inflammatory changes was observed, and the animals’ systemic statuses and behavioural changes were continuously monitored. The entire process was documented using imaging technology, including photographs and video recording, without affecting the survival of the animals.

### Murine Reproductive Model

#### Pairing and weighing

On day 30, each male was paired with one female from the same group and one healthy female control, with three mice in one cage, following a male-to-female ratio of 1:2. Animals were weighed on days 37, 41, 44, and 47.

#### Pregnancy and Delivery

On day 49 (19 days after pairing), all females were placed in individual cages and weighed daily. The pregnancy status of female mice was continuously observed until delivery. The survival of newborn mice was confirmed on the second day after birth. Observation of female mice lasted up to 21 days after separation.

#### Data Collection

Individual animal weights were measured, deliveries recorded, and neonatal mice counted (recording the number of alive pups and stillbirths).

#### The maximum body weight growth rate (MWGR)

MWGR is used to describe the physiological state of female mice during pregnancy and to detect the growth and development dynamics of the foetus. MWGR is the ratio of maximum body weight during pregnancy (usually the day before delivery in a normal pregnancy) to body weight at mating. When female mice become pregnant, their weight will gradually increase over the 1-2 weeks after conception. If the pregnancy is normal, weight gain will accelerate in the last week before delivery.

#### The average live birth rate (APR)

APR is an indicator of fertility, representing the average number of surviving pups born by each female mouse.

#### Humane Endpoint Implementation

When any abnormality occurred in the mice during the experiment or at the experiment’s conclusion, the mice were euthanised by asphyxiation through inhaling excessive carbon dioxide.

#### Database Establishment and Statistical Data Analysis

The original data of measurements and observations were entered into the database promptly. Analysis was performed based on raw data, and results were expressed as mean and standard deviation. Combined with the reproductive ability of male and female mice, statistical analysis was performed on the maximum weight growth rate (MWGR), alive pup rate (APR), and stillbirth rate using SPSS analysis software and methods. A P value of less than 0.05 was considered statistically significant. Results were analysed considering both statistical and biological significance.

## RESULTS

Influenced by clinical research references, when initially designing mouse reproductive models, we expected to observe indicators of reproductive abnormalities in infected animals, such as the proportion of neonatal mice born prematurely or with low birth weight. However, Balb/c mice, which are a multiple pregnancy mode animal, have a gestation period of 19-21 days, making the time standard for premature birth difficult to determine. Additionally, the more foetuses there are, the lower the average weight of the newborn mice. For example, twins to nonets are larger in weight and size at birth. Therefore, a newborn mouse being underweight doesn’t necessarily indicate abnormality.

In practice, survivor and stillbirth are direct consequences of pregnancy. However, the survival rate can more accurately reflect the pregnancy status because Balb/c mice have the habit of eating newborn mice considered abnormal, defective, or diseased, known as cannibalism (39). In our experience, most newborn pups can continue to grow if they survive more than 24 hours after birth. Therefore, the survival rate is used to evaluate the reproductive status, whilst the stillbirth rate can be used as an extra reference.

### Fertility analysis of normal healthy mice

In this experiment, animals in the healthy control group G7 and other groups were all 13-14 weeks old and in the sexual maturity stage at the time of mating. Each male was paired with two females. The results showed that all 16 females of G7 were pregnant, producing a total of 111 pups, of which 91 survived and 20 were stillborn. The average survival rate of females was 5.69±3.28, which meets the fertility standards of normal Balb/c mice. The average MWGR of healthy females was 170.1%±18.4% (Table 2).

**TABLE 2.** Fertility report-1, normal healthy and groups receiving single antibodies.

| Groups | Male serial number | pups per male |  |  | Female serial number | pups per female |  |  | MWGR |
| --- | --- | --- | --- | --- | --- | --- | --- | --- | --- |
|  |  | Survivors | Stillbirths | Subtotal |  | Survivors | Stillbirths | Subtotal |  |
| G7 Healthy Control | Ab-7-1 | 12 | 1 | 13 | G7-1-1 | 5 | 0 | 5 | 162.5% |
|  |  |  |  |  | G7-1-2 | 7 | 1 | 8 | 172.9% |
|  | Ab-7-2 | 16 | 1 | 17 | G7-2-1 | 11 | 1 | 12 | 191.8% |
|  |  |  |  |  | G7-2-2 | 5 | 0 | 5 | 154.6% |
|  | Ab-7-3 | 14 | 2 | 16 | G7-3-1 | 8 | 1 | 9 | 178.2% |
|  |  |  |  |  | G7-3-2 | 6 | 1 | 7 | 177.5% |
|  | Ab-7-4 | 13 | 4 | 17 | G7-4-1 | 6 | 3 | 9 | 179.0% |
|  |  |  |  |  | G7-4-2 | 7 | 1 | 8 | 175.5% |
|  | Ab-7-5 | 0 | 3 | 3 | G7-5-1 | 0 | 3 | 3 | 153.3% |
|  |  |  |  |  | G7-5-2 | 0 | 0 | 0 | 128.1% |
|  | Ab-7-6 | 17 | 1 | 18 | G7-6-1 | 9 | 0 | 9 | 178.2% |
|  |  |  |  |  | G7-6-2 | 8 | 1 | 9 | 183.8% |
|  | Ab-7-7 | 8 | 8 | 16 | G7-7-1 | 0 | 3 | 3 | 154.4% |
|  |  |  |  |  | G7-7-2 | 8 | 5 | 13 | 205.7% |
|  | Ab-7-8 | 11 | 0 | 11 | G7-8-1 | 7 | 0 | 7 | 176.6% |
|  |  |  |  |  | G7-8-2 | 4 | 0 | 4 | 154.7% |
| G1<br>1D2-2-1-3 | Ab-1-1 | 12 | 0 | 12 | G1-1 | 9 | 0 | 9 | 170.3% |
|  |  |  |  |  | G1C-1 | 3 | 0 | 3 | 142.3% |
|  | Ab-1-2 | 8 | 1 | 9 | G1-2 | 4 | 0 | 4 | 152.5% |
|  |  |  |  |  | G1C-2 | 4 | 1 | 5 | 151.4% |
|  | Ab-1-3 | 19 | 4 | 23 | G1-3 | 9 | 2 | 11 | 175.7% |
|  |  |  |  |  | G1C-3 | 10 | 2 | 12 | 197.4% |
|  | Ab-1-4 | 13 | 2 | 15 | G1-4 | 7 | 0 | 7 | 163.8% |
|  |  |  |  |  | G1C-4 | 6 | 2 | 8 | 171.6% |
|  | Ab-1-5 | 4 | 5 | 9 | G1-5 | 4 | 0 | 4 | 155.2% |
|  |  |  |  |  | G1C-5 | 0 | 5 | 5 | 147.8% |
|  | Ab-1-6 | 15 | 2 | 17 | G1-6 | 8 | 2 | 10 | 170.4% |
|  |  |  |  |  | G1C-6 | 7 | 0 | 7 | 171.6% |
|  | Ab-1-7 | 11 | 5 | 16 | G1-7 | 2 | 5 | 7 | 186.4% |
|  |  |  |  |  | G1C-7 | 9 | 0 | 9 | 179.2% |
|  | Ab-1-8 | 15 | 2 | 17 | G1-8 | 7 | 0 | 7 | 169.4% |
|  |  |  |  |  | G1C-8 | 8 | 2 | 10 | 168.5% |
| G2<br>1B11-4-4 | Ab-2-1 | 10 | 1 | 11 | G2-1 | 7 | 1 | 8 | 172.5% |
|  |  |  |  |  | G2C-1 | 3 | 0 | 3 | 144.8% |
|  | Ab-2-2 | 10 | 7 | 17 | G2-2 | 1 | 6 | 7 | 178.4% |
|  |  |  |  |  | G2C-2 | 9 | 1 | 10 | 168.6% |
|  | Ab-2-3 | 15 | 1 | 16 | G2-3 | 10 | 1 | 11 | 195.5% |
|  |  |  |  |  | G2C-3 | 5 | 0 | 5 | 179.7% |
|  | Ab-2-4 | 4 | 2 | 6 | G2-4 | 0 | 0 | 0 | 121.2% |
|  |  |  |  |  | G2C-4 | 4 | 2 | 6 | 154.6% |
|  | Ab-2-5 | 15 | 0 | 15 | G2-5 | 3 | 0 | 3 | 184.8% |
|  |  |  |  |  | G2C-5 | 12 | 0 | 12 | 171.3% |
|  | Ab-2-6 | 0 | 0 | 0 | G2-6 | 0 | 0 | 0 | 110.4% |
|  |  |  |  |  | G2C-6 | 0 | 0 | 0 | 117.1% |
|  | Ab-2-7 | 10 | 1 | 11 | G2-7 | 3 | 0 | 3 | 159.4% |
|  |  |  |  |  | G2C-7 | 7 | 1 | 8 | 187.9% |
|  | Ab-2-8 | 14 | 2 | 16 | G2-8 | 8 | 2 | 10 | 198.1% |
|  |  |  |  |  | G2C-8 | 6 | 0 | 6 | 163.4% |
| G3<br>1C3-7-8 | Ab-3-1 | 12 | 0 | 12 | G3-1 | 4 | 0 | 4 | 165.3% |
|  |  |  |  |  | G3C-1 | 8 | 0 | 8 | 174.4% |
|  | Ab-3-2 | 8 | 2 | 10 | G3-2 | 2 | 0 | 2 | 155.4% |
|  |  |  |  |  | G3C-2 | 6 | 2 | 8 | 154.5% |
|  | Ab-3-3 | 13 | 3 | 16 | G3-3 | 6 | 1 | 7 | 161.2% |
|  |  |  |  |  | G3C-3 | 7 | 2 | 9 | 179.9% |
|  | Ab-3-4 | 14 | 0 | 14 | G3-4 | 4 | 0 | 4 | 161.9% |
|  |  |  |  |  | G3C-4 | 10 | 0 | 10 | 193.6% |
|  | Ab-3-5 | 7 | 1 | 8 | G3-5 | 7 | 1 | 8 | 175.3% |
|  |  |  |  |  | G3C-5 | 0 | 0 | 0 | 121.0% |
|  | Ab-3-6 | 11 | 4 | 15 | G3-6 | 6 | 0 | 6 | 166.7% |
|  |  |  |  |  | G3C-6 | 5 | 4 | 9 | 182.2% |
|  | Ab-3-7 | 9 | 0 | 9 | G3-7 | 0 | 0 | 0 | 111.0% |
|  |  |  |  |  | G3C-7 | 9 | 0 | 9 | 170.0% |
|  | Ab-3-8 | 18 | 2 | 20 | G3-8 | 10 | 1 | 11 | 171.6% |
|  |  |  |  |  | G3C-8 | 8 | 1 | 9 | 173.5% |

Three cases of adverse pregnancy (APO) occurred in this group. Females G7-5-1 and G7-5-2, mated with male Ab-7-5, failed to produce any healthy and living offspring. This may indicate that the male mouse Ab-7-5 is more likely responsible for APO rather than the two females. Female G7-7-1 produced only three stillbirths.

The healthy control group serves as the standard for normal reproduction in this experiment. Clinically, substandard sperm quality can lead to infertility, while adverse pregnancy may result in stillbirth or premature offspring. Our experimental results suggest that in natural reproduction, the male’s condition may also have an effect on adverse pregnancy outcomes, including stillbirth.

### Fertility analysis of mice infected with *P. gingivalis*

In infection group G4, both males and females were subcutaneously challenged with *P. gingivalis* on the 6th and 22nd days after grouping. On day 30, each infected male was paired with an infected female and a healthy control female. The purpose of adding normal healthy females was to observe the effect of infected males on the fertility of both normal and infected female mice. The average survival rate (APR) of G4 pups was 4.5±4.07, and the average of MWGR was 162.1%±23.6% (Table 3). A total of 90 pups were born from 16 female mice, of which 72 survived and 18 were stillborn.

**TABLE 3.** Fertility report-2, infection mice and groups receiving mixture of antibodies.

| Groups | Male serial number | pups per male |  |  | Female serial number | pups per female |  |  | MWGR |
| --- | --- | --- | --- | --- | --- | --- | --- | --- | --- |
|  |  | Survivors | Stillbirths | Subtotal |  | Survivors | Stillbirths | Subtotal |  |
| G4 Infection Control | Ab-4-1 | 10 | 4 | 14 | G4-1 | 6 | 1 | 7 | 173.3% |
|  |  |  |  |  | G4C-1 | 4 | 3 | 7 | 175.8% |
|  | Ab-4-2 | 23 | 2 | 25 | G4-2 | 10 | 0 | 10 | 185.3% |
|  |  |  |  |  | G4C-2 | 13 | 2 | 15 | 201.0% |
|  | Ab-4-3 | 10 | 3 | 13 | G4-3 | 5 | 1 | 6 | 166.9% |
|  |  |  |  |  | G4C-3 | 5 | 2 | 7 | 162.3% |
|  | Ab-4-4 | 0 | 5 | 5 | G4-4 | 0 | 5 | 5 | 162.3% |
|  |  |  |  |  | G4C-4 | 0 | 0 | 0 | 115.1% |
|  | Ab-4-5 | 10 | 0 | 10 | G4-5 | 0 | 0 | 0 | 138.9% |
|  |  |  |  |  | G4C-5 | 10 | 0 | 10 | 181.1% |
|  | Ab-4-6 | 7 | 4 | 11 | G4-6 | 6 | 1 | 7 | 185.4% |
|  |  |  |  |  | G4C-6 | 1 | 3 | 4 | 171.1% |
|  | Ab-4-7 | 10 | 0 | 10 | G4-7 | 4 | 0 | 4 | 156.6% |
|  |  |  |  |  | G4C-7 | 6 | 0 | 6 | 160.3% |
|  | Ab-4-8 | 1 | 1 | 2 | G4-8 | 1 | 1 | 2 | 133.9% |
|  |  |  |  |  | G4C-8 | 0 | 0 | 0 | 124.7% |
| G5 Mabs 1B11-4-4 and 1C3-7-8 | Ab-5-1 | 5 | 0 | 5 | G5-1 | 0 | 0 | 0 | 112.8% |
|  |  |  |  |  | G5C-1 | 5 | 0 | 5 | 160.2% |
|  | Ab-5-2 | 11 | 1 | 12 | G5-2 | 3 | 1 | 4 | 157.8% |
|  |  |  |  |  | G5C-2 | 8 | 0 | 8 | 175.7% |
|  | Ab-5-3 | 7 | 2 | 9 | G5-3 | 7 | 1 | 8 | 182.3% |
|  |  |  |  |  | G5C-3 | 0 | 1 | 1 | 152.9% |
|  | Ab-5-4 | 13 | 1 | 14 | G5-4 | 2 | 0 | 2 | 148.5% |
|  |  |  |  |  | G5C-4 | 11 | 1 | 12 | 200.6% |
|  | Ab-5-5 | 6 | 0 | 6 | G5-5 | 6 | 0 | 6 | 173.0% |
|  |  |  |  |  | G5C-5 | 0 | 0 | 0 | 117.1% |
|  | Ab-5-6 | 9 | 0 | 9 | G5-6 | 0 | 0 | 0 | 133.0% |
|  |  |  |  |  | G5C-6 | 9 | 0 | 9 | 175.7% |
|  | Ab-5-7 | 0 | 2 | 2 | G5-7 | 0 | 2 | 2 | 146.9% |
|  |  |  |  |  | G5C-7 | 0 | 0 | 0 | 107.9% |
|  | Ab-5-8 | 12 | 2 | 14 | G5-8 | 8 | 0 | 8 | 183.3% |
|  |  |  |  |  | G5C-8 | 4 | 2 | 6 | 161.3% |
| G6 Mabs 1B11-4-4 and 1D2-2-1-3 | Ab-6-1 | 10 | 6 | 16 | G6-1 | 7 | 2 | 9 | 188.1% |
|  |  |  |  |  | G6C-1 | 3 | 4 | 7 | 159.2% |
|  | Ab-6-2 | 7 | 0 | 7 | G6-2 | 0 | 0 | 0 | 120.8% |
|  |  |  |  |  | G6C-2 | 7 | 0 | 7 | 178.1% |
|  | Ab-6-3 | 0 | 0 | 0 | G6-3 | 0 | 0 | 0 | 147.0% |
|  |  |  |  |  | G6C-3 | 0 | 0 | 0 | 119.4% |
|  | Ab-6-4 | 10 | 3 | 13 | G6-4 | 5 | 2 | 7 | 188.0% |
|  |  |  |  |  | G6C-4 | 5 | 1 | 6 | 162.7% |
|  | Ab-6-5 | 5 | 1 | 6 | G6-5 | 0 | 0 | 0 | 112.2% |
|  |  |  |  |  | G6C-5 | 5 | 1 | 6 | 176.6% |
|  | Ab-6-6 | 0 | 1 | 1 | G6-6 | 0 | 0 | 0 | 121.6% |
|  |  |  |  |  | G6C-6 | 0 | 1 | 1 | 133.6% |
|  | Ab-6-7 | 4 | 1 | 5 | G6-7 | 4 | 0 | 4 | 159.1% |
|  |  |  |  |  | G6C-7 | 0 | 1 | 1 | 131.8% |
|  | Ab-6-8 | 6 | 0 | 6 | G6-8 | 4 | 0 | 4 | 161.3% |
|  |  |  |  |  | G6C-8 | 2 | 0 | 2 | 156.4% |

Fertility assessment: Male Ab-4-4 was paired with G4-4 and G4C-4, and both females failed to produce healthy offsprings. Male Ab-4-8 was paired with two females, resulting in infertility in G4C-8 and G4-8 producing one live pup and one stillbirth. Other adverse pregnancies occurred randomly in different groups, such as female G4C-1, G4-5 and G4C-6.

The results for the G4 showed interesting polarity. Male Ab-4-2 had high reproductive success, producing the highest number of pups and survivors throughout the experiment. Two females paired with this male gave birth to a total of 25 pups, with 2 stillbirths. Notably, the conception time for G4-2 and G4C-2 occurred 10 days after mating, suggesting a second pregnancy after early failure of the first. Typically, first-time pregnant mice produce fewer pups than multiparous mice.

Compared with the healthy control group G7, even taking into account the success of Ab-4-2, the pregnancy APR and MWGR of the G4 group were lower. Although local subcutaneous invasion of bacteria and female reproductive organs involve different anatomical tissues, our data show that *P. gingivalis* infection is associated with adverse pregnancy outcomes, suggesting that the bacterium or its components circulate systemically.

Experimental results showed that when infected males were mated with both infected and healthy control females, the average MWGR and APR of infected females were 4.43±3.51 and 162.8%±19.2%, respectively, and those of the control group were 4.88±4.73 and 161.4%±28.7%. Statistical analysis showed no significant difference, between the two female groups, further suggesting that *P. gingivalis* infection in females does not significantly affect reproduction, whereas male infection status appears to be the major factor leading to reproductive abnormalities.

### Fertility analysis of mice receiving individual monoclonal antibodies

Three different monoclonal antibody drugs were used. The injection dose was 100μg per mouse, with two intraperitoneal injections 4 days apart before *P. gingivalis* challenge. During the recovery period from infection (14 days after the first challenge), mice were reinjected with antibodies (100μg) followed by a second *P. gingivalis* challenge. The purpose was to evaluate the passive immune protective function of these antibodies.

Group G1 were given monoclonal antibody 1D2-2-1-3. This antibody was raised after immunisation with *P. gingivalis* outer membrane protein RagB-3. Monoclonal antibodies 1B11-4-4 and 1C3-7-8 used in groups G2 and G3 were prepared after immunisation with *P. gingivalis* outer membrane protein RagB-4 (11).

Results showed that animals in groups G1-G3 demonstrated positive and effective protective functions in the reproductive model. The fertility indicators of females in these groups were similar to the normal control group G7 and better than the infection group G4. The average MWGR were 167.1%±14.6%, 163.0% ±27.2%, and 163.6%±21.1%, and APR 6.06±2.91, 4.88±3.76, and 5.75±3.13, respectively (Tables 2). Paired data statistical analysis showed no significant difference between females receiving antibodies and *P. gingivalis* infection, and healthy control females. The results indicate that the three monoclonal antibodies provided immune protection after administration, suggesting that circulating antibodies can inhibit local bacterial infection and prevent reproductive abnormalities induced by the bacterial infection.

### Fertility analysis of mice receiving mixed monoclonal antibodies

Combination use of monoclonal antibodies was proposed to provide better protection. However, we observed a higher rate of adverse pregnancy outcomes in these groups compared to single antibody use alone.

Group G5 animals, which received mixed monoclonal antibodies 1C3-7-8 and 1B11-4-4, had MWGR and APR of 155.6%±27.0% and 3.94±3.84, respectively. Sixteen females produced 71 pups, of which 63 survived and 8 were stillborn. Six females failed to produce live pups (Table 3).

Group G6 animals, which received mixed monoclonal antibodies 1D2-2-1-3 and 1B11-4-4, had MWGR and APR of 151.0%±25.2% and 2.63±2.68, respectively. Sixteen female mice gave birth to a total of 54 pups, of which 42 survived and 12 were stillborn. Seven females failed to produce live pups (Table 3).

Animals in groups G5 and G6 received 100μg in total of two antibodies mixed in equal amounts, with each antibody half the dose of the same antibody drug used in groups G1-G3. Receiving mixed antibodies and bacterial infection caused more severe reproductive damage than bacterial infection alone, whereas single antibody treatment alone was immunoprotective. The APR values of the mixed antibody groups G5 and G6 were significantly lower than those of the single antibody treatment groups G1-G3 and the healthy control group G7, with survival rates only 46-70% of the healthy control group. This suggests that mixed antibodies circulating in the body encounter P. *gingivalis*, damaging tissue cells and affecting reproduction.

The mixture of multiple antibodies appeared to cause serious reproductive damage. We temporarily named this phenomenon the “antibody pool effect”. Based on our experimental data and results, we proposed a theory for the pathogenic mechanism by which *P. gingivalis* causes reproductive damage through immune complexes formed by the combination of antigens and multivalent antibodies.

## SUMMARY OF MOUSE REPRODUCTIVE MODEL

A total of 56 male mice and 112 female mice were utilised in this experiment, giving birth to a total of 640 pups, including 534 live pups and 106 stillbirths (Table 4). We used SPSS generalised linear models to analyse the collected data and specifically used generalised estimating equations (GEE) to process and compare each group of data. Table 5 lists the values extrapolated from APR raw data, and Table 6 lists the pairwise comparison results between each set of data. The results showed that the APR values of the G6 group were significantly worse than those of the G1, G2, G3, and G7 groups. The differences between the G1, G3, and G5 groups were also statistically significant (Fig. 1). This method was also used to analyse body weight growth MWGR data in female mice, showing significant differences between G7 and G5/G6, and G1 and G6 (Table 7-8 and Fig. 2).

**TABLE 4.** Summary of information on total number of pups.

| Groups | Survivors |  |  | Stillbirths |  |  | Total pups |  |  |
| --- | --- | --- | --- | --- | --- | --- | --- | --- | --- |
|  | Number | Average | Standard Deviation | Number | Average | Standard Deviation | Number | Average | STD |
| G1 | 97 | 6.06 | 2.91 | 21 | 1.31 | 1.70 | 118 | 7.38 | 2.68 |
| G2 | 78 | 4.88 | 3.76 | 14 | 0.88 | 1.54 | 92 | 5.75 | 3.97 |
| G3 | 92 | 5.75 | 3.13 | 12 | 0.75 | 1.13 | 104 | 6.50 | 3.48 |
| G4 | 71 | 4.44 | 4.05 | 19 | 1.19 | 1.47 | 90 | 5.63 | 4.06 |
| G5 | 63 | 3.94 | 3.84 | 8 | 0.50 | 0.73 | 71 | 4.44 | 3.86 |
| G6 | 42 | 2.63 | 2.68 | 12 | 0.75 | 1.13 | 54 | 3.38 | 3.22 |
| G7 | 91 | 5.69 | 3.28 | 20 | 1.25 | 1.48 | 111 | 6.94 | 3.43 |
| Total | 534 |  |  | 106 |  |  | 640 |  |  |

**TABLE 5.** Values related to survivals estimated from the generalized estimating equation GEE.

| Groups | Average | Standard Deviation | 95% Wald Confidence Interval |  |
| --- | --- | --- | --- | --- |
|  |  |  | Lower limit | Upper limit |
| G1 | 6.06 | 0.76 | 4.57 | 7.56 |
| G2 | 4.88 | 0.89 | 3.14 | 6.61 |
| G3 | 5.75 | 0.59 | 4.59 | 6.91 |
| G4 | 4.44 | 1.17 | 2.20 | 6.80 |
| G5 | 3.94 | 0.71 | 2.55 | 5.33 |
| G6 | 2.62 | 0.64 | 1.37 | 3.88 |
| G7 | 5.69 | 0.89 | 3.94 | 7.44 |

**TABLE 6.**
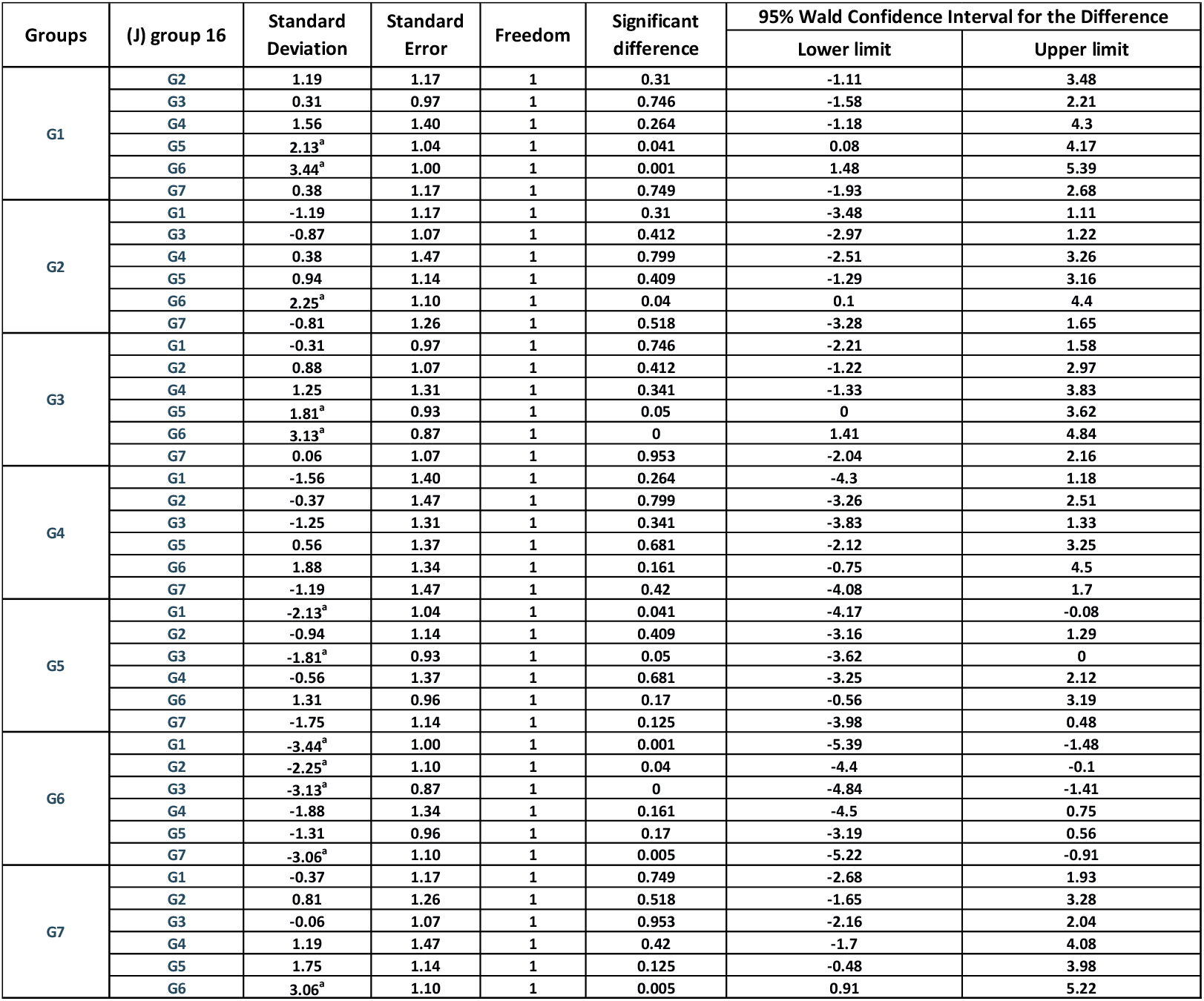
APR multiple comparisons (generalized estimating equation GEE)

**TABLE 7.**
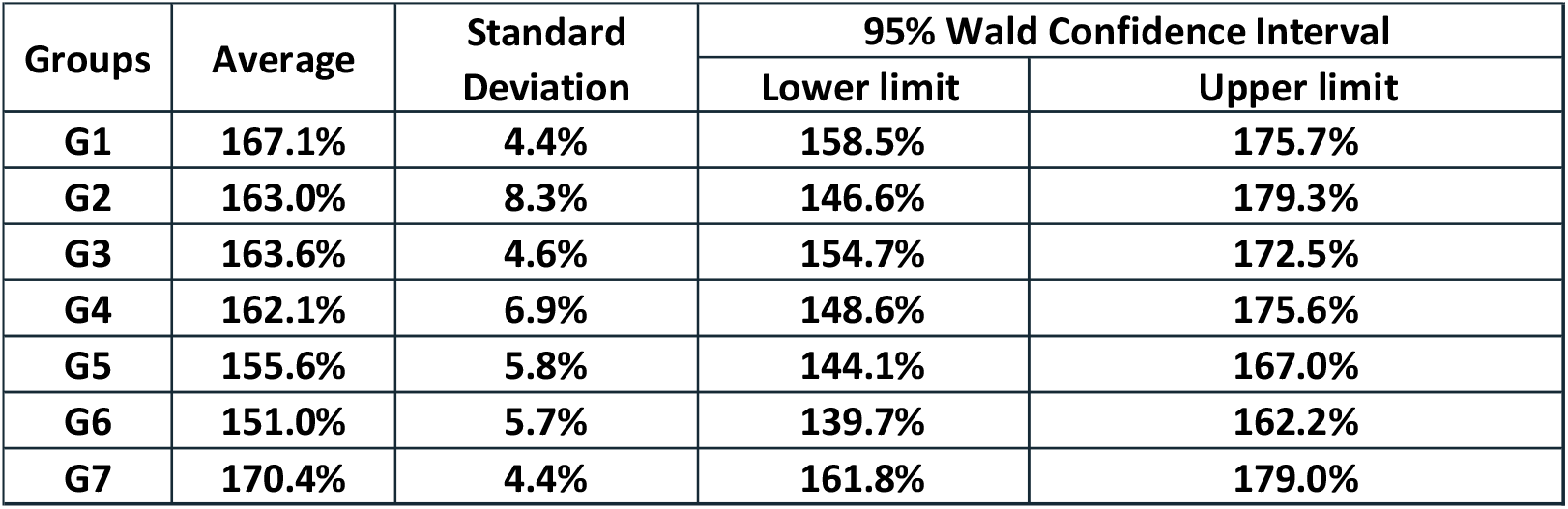
Values related to MWGR estimated from the generalized estimating equation GEE.

**TABLE 8.**
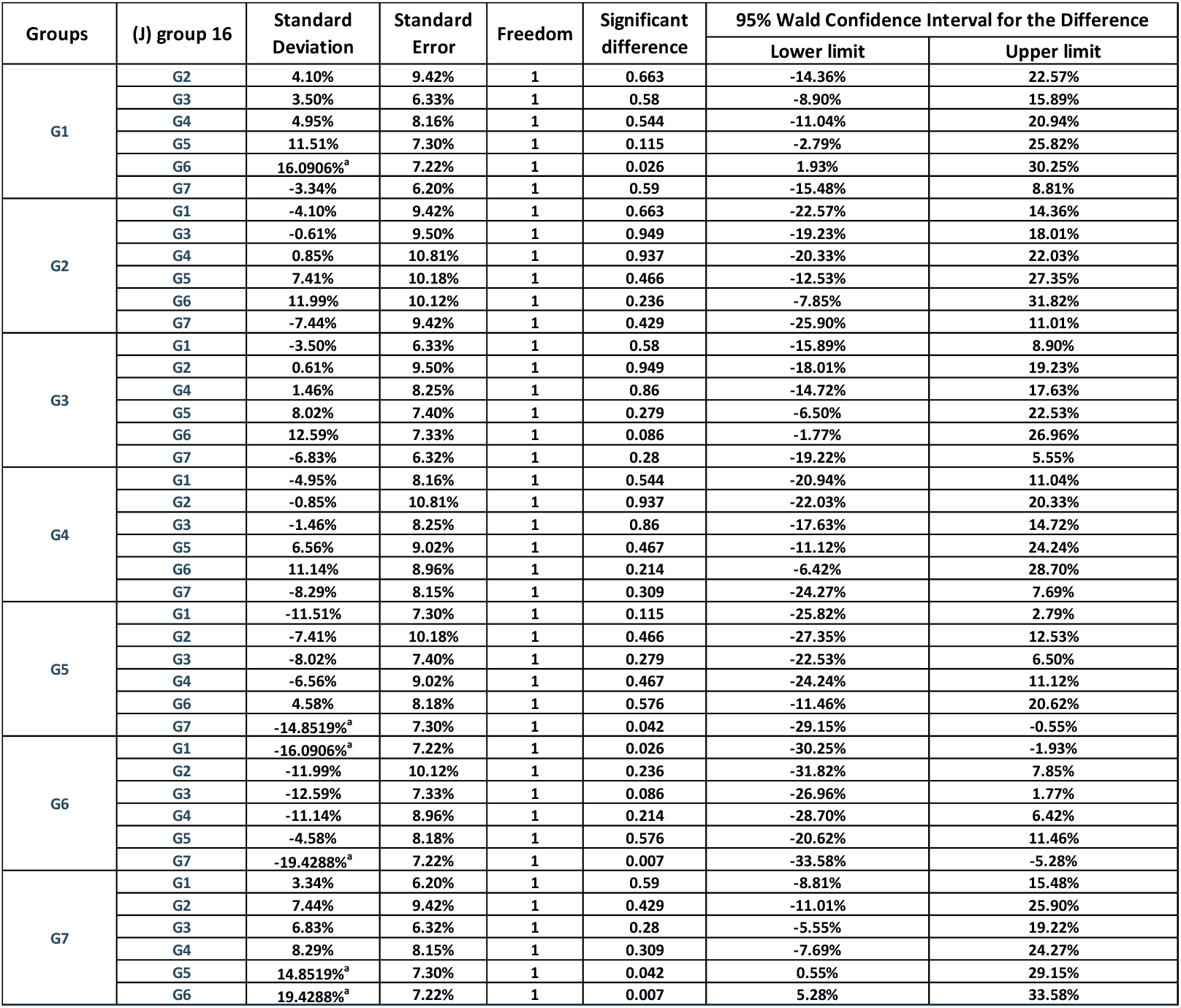
MWGR multiple comparisons (generalized estimating equation GEE)

| Groups | (J) group 16 | Standard Deviation | Standard Error | Freedom | Significant difference | 95% Wald Confidence Interval for the Difference |  |
| --- | --- | --- | --- | --- | --- | --- | --- |
|  |  |  |  |  |  | Lower limit | Upper limit |
| G1 | G2 | 4.10% | 9.42% | 1 | 0.663 | -14.36% | 22.57% |
|  | G3 | 3.50% | 6.33% | 1 | 0.58 | -8.90% | 15.89% |
|  | G4 | 4.95% | 8.16% | 1 | 0.544 | -11.04% | 20.94% |
|  | G5 | 11.51% | 7.30% | 1 | 0.115 | -2.79% | 25.82% |
|  | G6 | 16.0906%<sup>a</sup> | 7.22% | 1 | 0.026 | 1.93% | 30.25% |
|  | G7 | -3.34% | 6.20% | 1 | 0.59 | -15.48% | 8.81% |
| G2 | G1 | -4.10% | 9.42% | 1 | 0.663 | -22.57% | 14.36% |
|  | G3 | -0.61% | 9.50% | 1 | 0.949 | -19.23% | 18.01% |
|  | G4 | 0.85% | 10.81% | 1 | 0.937 | -20.33% | 22.03% |
|  | G5 | 7.41% | 10.18% | 1 | 0.466 | -12.53% | 27.35% |
|  | G6 | 11.99% | 10.12% | 1 | 0.236 | -7.85% | 31.82% |
|  | G7 | -7.44% | 9.42% | 1 | 0.429 | -25.90% | 11.01% |
| G3 | G1 | -3.50% | 6.33% | 1 | 0.58 | -15.89% | 8.90% |
|  | G2 | 0.61% | 9.50% | 1 | 0.949 | -18.01% | 19.23% |
|  | G4 | 1.46% | 8.25% | 1 | 0.86 | -14.72% | 17.63% |
|  | G5 | 8.02% | 7.40% | 1 | 0.279 | -6.50% | 22.53% |
|  | G6 | 12.59% | 7.33% | 1 | 0.086 | -1.77% | 26.96% |
|  | G7 | -6.83% | 6.32% | 1 | 0.28 | -19.22% | 5.55% |
| G4 | G1 | -4.95% | 8.16% | 1 | 0.544 | -20.94% | 11.04% |
|  | G2 | -0.85% | 10.81% | 1 | 0.937 | -22.03% | 20.33% |
|  | G3 | -1.46% | 8.25% | 1 | 0.86 | -17.63% | 14.72% |
|  | G5 | 6.56% | 9.02% | 1 | 0.467 | -11.12% | 24.24% |
|  | G6 | 11.14% | 8.96% | 1 | 0.214 | -6.42% | 28.70% |
|  | G7 | -8.29% | 8.15% | 1 | 0.309 | -24.27% | 7.69% |
| G5 | G1 | -11.51% | 7.30% | 1 | 0.115 | -25.82% | 2.79% |
|  | G2 | -7.41% | 10.18% | 1 | 0.466 | -27.35% | 12.53% |
|  | G3 | -8.02% | 7.40% | 1 | 0.279 | -22.53% | 6.50% |
|  | G4 | -6.56% | 9.02% | 1 | 0.467 | -24.24% | 11.12% |
|  | G6 | 4.58% | 8.18% | 1 | 0.576 | -11.46% | 20.62% |
|  | G7 | -14.8519%<sup>a</sup> | 7.30% | 1 | 0.042 | -29.15% | -0.55% |
| G6 | G1 | -16.0906%<sup>a</sup> | 7.22% | 1 | 0.026 | -30.25% | -1.93% |
|  | G2 | -11.99% | 10.12% | 1 | 0.236 | -31.82% | 7.85% |
|  | G3 | -12.59% | 7.33% | 1 | 0.086 | -26.96% | 1.77% |
|  | G4 | -11.14% | 8.96% | 1 | 0.214 | -28.70% | 6.42% |
|  | G5 | -4.58% | 8.18% | 1 | 0.576 | -20.62% | 11.46% |
|  | G7 | -19.4288%<sup>a</sup> | 7.22% | 1 | 0.007 | -33.58% | -5.28% |
| G7 | G1 | 3.34% | 6.20% | 1 | 0.59 | -8.81% | 15.48% |
|  | G2 | 7.44% | 9.42% | 1 | 0.429 | -11.01% | 25.90% |
|  | G3 | 6.83% | 6.32% | 1 | 0.28 | -5.55% | 19.22% |
|  | G4 | 8.29% | 8.15% | 1 | 0.309 | -7.69% | 24.27% |
|  | G5 | 14.8519%<sup>a</sup> | 7.30% | 1 | 0.042 | 0.55% | 29.15% |
|  | G6 | 19.4288%<sup>a</sup> | 7.22% | 1 | 0.007 | 5.28% | 33.58% |

**FIG 1.**
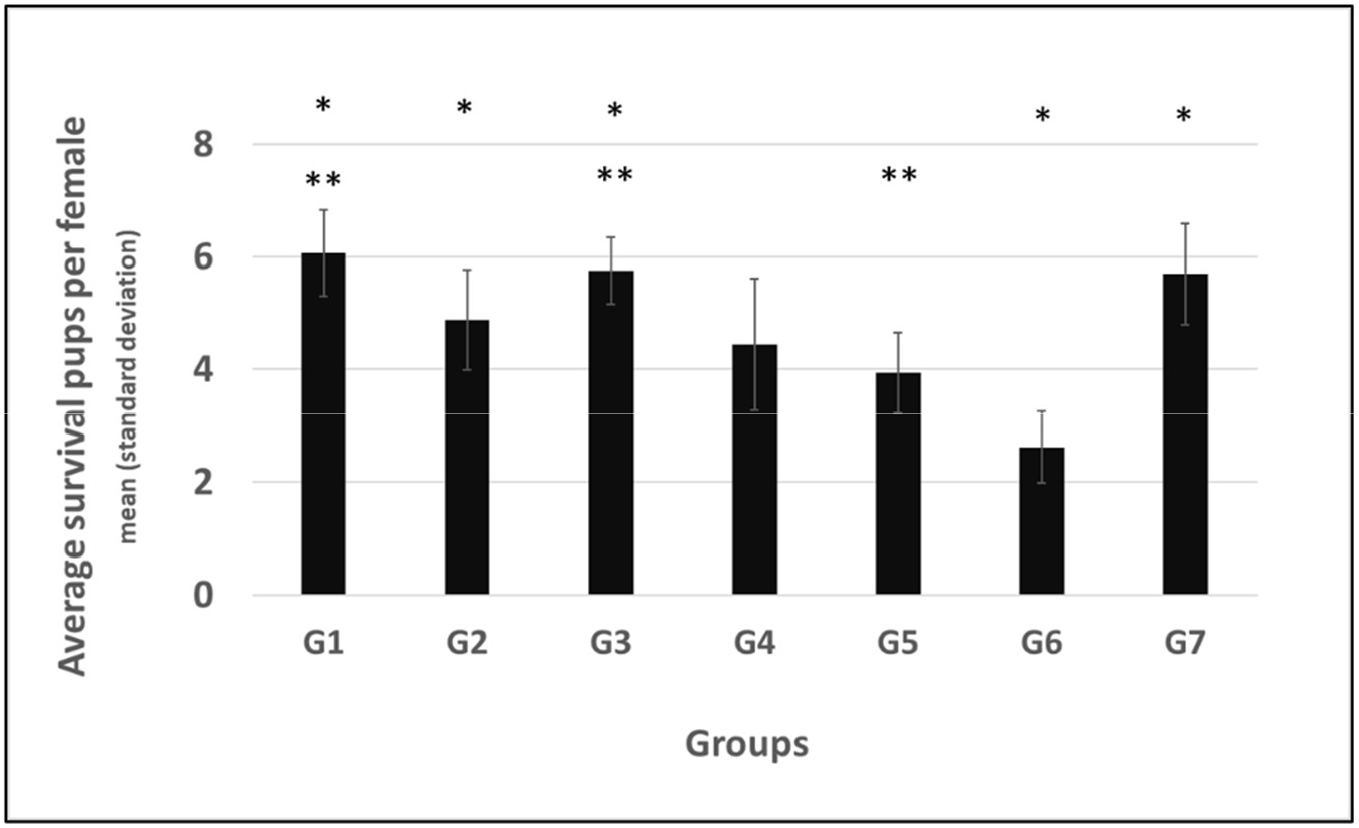
Effects of *P. gingivalis* infection on mouse reproduction (APR) *The survival rates of offspring in group G6 (receiving a mixture of monoclonal antibodies) were significantly worse than in groups G1, G2, G3 (receiving a single monoclonal antibody) and G7 (healthy controls) (P<0.05); ** The survival rates of offspring in group G5 (receiving a mixture of monoclonal antibodies) were significantly worse than in groups G1 and G3 (receiving a single monoclonal antibody) (P<0.05).

**FIG 2.**
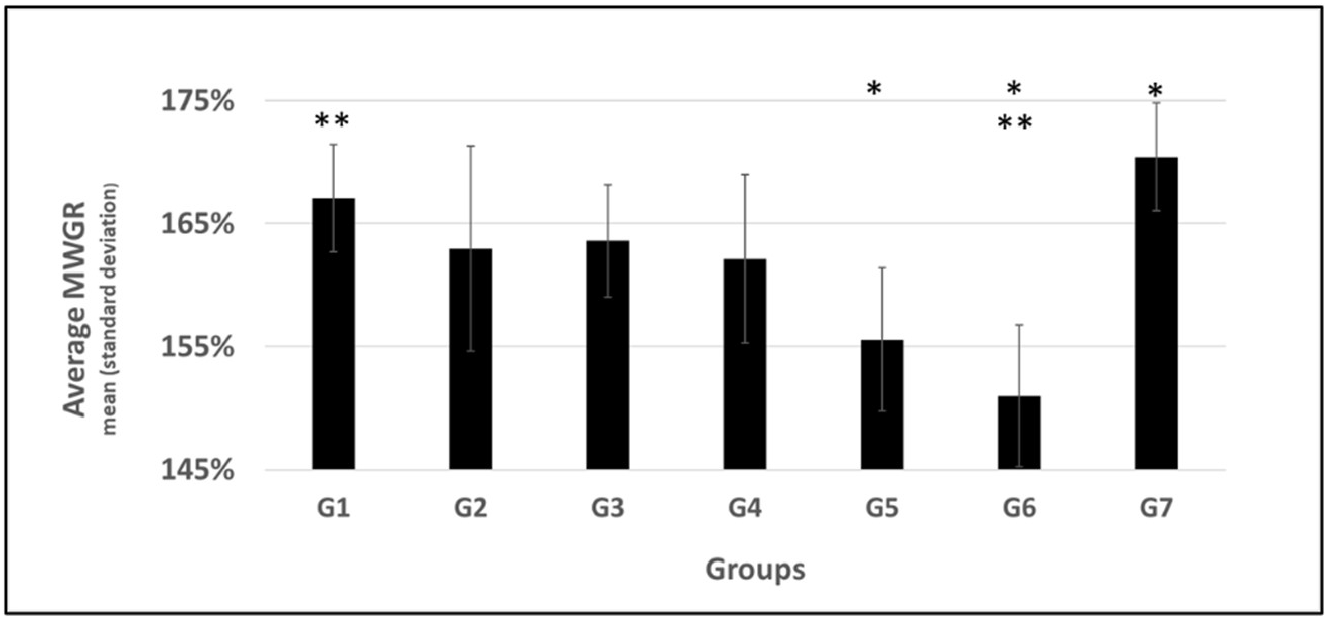
Effects of *P. gingivalis* infection on mouse reproduction (MWGR) *Female maximum weight growth rate (MWGR) in groups G5 and G6 (receiving a mixture of monoclonal antibodies) were significantly worse than in group G7 (healthy controls) (P<0.05); ** The MWGR in group G6 (receiving a mixture of monoclonal antibodies) were significantly worse than in group G1 (receiving a single monoclonal antibody) (P<0.05).

Each group from G1 to G6 included two sets of female mice: 8 infected and 8 healthy. SPSS statistical paired analysis did not reveal any significant differences in the data within the groups (Tables 9). The results suggest that *P. gingivalis* infection in females does not significantly affect the physiological state and reproduction of pregnant mice, while male infection status appears to play a leading role in reproductive abnormalities.

**TABLE 9.** Paired samples SPSS tests.

| APR Paired samples test |  | Paired mean difference | Standard Deviation | Standard error of the mean | Difference 95% confidence interval |  | T | Freedom | Significance (two-tailed) |
| --- | --- | --- | --- | --- | --- | --- | --- | --- | --- |
|  |  |  |  |  | Lower limit | Upper limit |  |  |  |
| Pair 1 | G1-control vs G1-antibody | -0.375 | 3.852 | 1.362 | -3.596 | 2.846 | -0.275 | 7 | 0.791 |
| Pair 2 | G2-control vs G2-antibody | 1.75 | 5.312 | 1.878 | -2.691 | 6.191 | 0.932 | 7 | 0.382 |
| Pair 3 | G3-control vs G3-antibody | 1.75 | 5.064 | 1.79 | -2.484 | 5.984 | 0.977 | 7 | 0.361 |
| Pair 4 | G4-control vs G4-antibody | 0.875 | 4.422 | 1.563 | -2.822 | 4.572 | 0.56 | 7 | 0.593 |
| Pair 5 | G5-control vs G5-antibody | 1.375 | 6.523 | 2.306 | -4.079 | 6.829 | 0.596 | 7 | 0.57 |
| Pair 6 | G6-control vs G6-antibody | 0.25 | 3.955 | 1.398 | -3.057 | 3.557 | 0.179 | 7 | 0.863 |
| MWGR Paired samples test |  |  |  |  |  |  |  |  |  |
| Pair 1 | G1-control vs G1-antibody | -1.74% | 14.15% | 5.00% | -13.57% | 10.09% | -0.347 | 7 | 0.738 |
| Pair 2 | G2-control vs G2-antibody | -4.09% | 24.87% | 8.79% | -24.88% | 16.71% | -0.465 | 7 | 0.656 |
| Pair 3 | G3-control vs G3-antibody | 10.07% | 32.28% | 11.41% | -16.91% | 37.06% | 0.883 | 7 | 0.407 |
| Pair 4 | G4-control vs G4-antibody | -1.40% | 25.57% | 9.04% | -22.78% | 19.98% | -0.155 | 7 | 0.881 |
| Pair 5 | G5-control vs G5-antibody | 1.71% | 43.24% | 15.29% | -34.44% | 37.86% | 0.112 | 7 | 0.914 |
| Pair 6 | G6-control vs G6-antibody | 2.46% | 38.78% | 13.71% | -29.97% | 34.88% | 0.179 | 7 | 0.863 |

Groups G5 and G6, which received mixed antibodies followed by *P. gingivalis* infection, had worse reproductive outcomes than the infection-only group G4. This suggests that a mixed antibody combination may have been causative. Group G6 had the most severe reproductive damage of all study groups, consistent with the data of patent PCT/EP2022/075842 (11). This indicates that multiple antibodies present in the body may cause damage to an animal’s reproductive system when encountering *P. gingivalis*.

## DISCUSSION AND CONCLUSION

Our preliminary clinical serological survey results showed a high detection rate of antibodies against *P. gingivalis* outer membrane proteins in the sera of patients with cardiovascular and geriatric diseases (11). There are many reports in the literature regarding the detection of antibodies to oral bacteria in the sera of patients with periodontal disease, cardiovascular disease, and Alzheimer’s disease (40-54). The presence of anti-*P. gingivalis* antibodies in patient sera suggests that these subjects either had natural antibodies or a history of *P. gingivalis* infection. The presence of chronic *P. gingivalis* infection or corresponding chronic tissue damage indicates an ongoing adjustment in the physiological and pathological balance between infection and immune resistance.

While there is an immune response in infected patients, the immune system does not promptly or effectively clear the infection. This persistent infection may lead to repeated cycles of chronic tissue damage and immune response, highlighting a need for improved therapeutic strategies to enhance the immune system’s ability to eliminate *P. gingivalis* effectively (55).

### Reproductive Model Insights

Our study demonstrated that the infection status of male mice played a critical role in reproductive outcomes. The significant reduction in reproductive metrics such as the Average Pup Rate (APR) and Maximum Weight Growth Rate (MWGR) in infected male mice underscores the substantial impact of *P. gingivalis* on male fertility. This effect was seen irrespective of whether the female mice were infected or healthy, suggesting that male infection alone can severely impair reproductive success.

Interestingly, while female infection did not significantly affect reproductive outcomes, the mixture of multiple monoclonal antibodies administered to both male and female mice led to severe reproductive damage, a phenomenon we termed the “antibody pool effect.” This adverse outcome may be due to the formation of immune complexes that exacerbate tissue damage rather than providing the intended protective effect.

### Implications for Human Health

The implications of these findings for human health are profound. Given the high prevalence of *P. gingivalis* antibodies in patients with chronic diseases, it is plausible that similar mechanisms of immune complex formation and chronic tissue damage may occur in humans. This underscores the potential risks associated with chronic *P. gingivalis* infection, not only in terms of direct tissue damage but also through complex immune-mediated mechanisms that may exacerbate disease progression.

### Future Directions

Future research should focus on understanding the precise mechanisms by which *P. gingivalis* and its antigens interact with the host immune system to cause chronic damage. Therapeutic strategies should aim to enhance the clearance of *P. gingivalis* while avoiding the formation of harmful immune complexes. Additionally, the development of targeted immunotherapies that can selectively enhance the immune response to *P. gingivalis* without triggering adverse effects will be crucial.

Overall, our study highlights the intricate balance between infection, immune response, and tissue damage, emphasising the need for a nuanced approach to treating chronic bacterial infections and their potential systemic impacts.

## ACKNOWLEDGMENTS

We thank Professors Mike Curtis (King’s College, UK), Mark Roberts (University of Glasgow, UK), Liwei Lv (University of Hong Kong, CN), and Dr. Tian Xu (Nanjing Forestry University, CN) for their helpful discussions and suggestions during manuscript preparation. We also extend our gratitude to Professors Junrong Li (Jiangsu University, CN) and Duolao Wang (University of Liverpool, UK) for their support in the statistical analysis of data. We thank the R&D platforms HuaBio (Hangzhou, CN) and Hongren Biopharm (Wuhan, CN) for their collaboration. We would like to thank ChatGPT, a language model developed by OpenAI, for assistance in the writing process of this paper.

